# Deciphering the Role of Acetate in Metabolic Adaptation and Drug Resistance in Non-Small Cell Lung Cancer

**DOI:** 10.1101/2025.07.04.663159

**Authors:** Giorgia Maroni, Eva Cabrera San Millan, Beatrice Campanella, Massimo Onor, Giovanni Cercignani, Beatrice Muscatello, Giulia Braccini, Raffaella Mercatelli, Alice Chiodi, Ettore Mosca, Elena Levantini, Emilia Bramanti

## Abstract

Resistance to targeted therapies remains a major challenge in EGFR-mutant non-small cell lung cancer (NSCLC). Here, we describe a novel metabolic adaptation in osimertinib-resistant cells characterized by elevated acetate levels and activation of an unconventional pyruvate–acetaldehyde–acetate (PAA) shunt. Integrated transcriptomic, exometabolomic, and functional analyses reveal suppression of canonical metabolic pathways and upregulation of ALDH2 and ALDH7A1, which mediate the NADP^+^-dependent oxidation of acetaldehyde to acetate, generating NADPH. This shift supports reducing power essential for biosynthesis and redox balance under conditions of oxidative pentose phosphate inhibition.

These metabolic changes promote endurance in resistant cells and rewire the interplay between glycolysis, the pentose phosphate pathway, and the tricarboxylic acid cycle, offering a *de novo* bypass for anaplerosis and bioenergetics. Systematic metabolite profiling revealed distinct transcriptomic and metabolic signatures distinguishing resistant from parental cells.

Together, these findings depict a unique, resistance-driven adaptive metabolic shift, and uncover potential therapeutic vulnerabilities in osimertinib-resistant NSCLC.

## 1. Introduction

Mitochondrial oxidative phosphorylation (OXPHOS), localized to the inner mitochondrial membrane, is the primary source of ATP in most eukaryotic cells. It couples electron flow along the respiratory chain [electron transport chain (ETC)] with ATP synthesis via ATP synthase, yielding ∼30-34 ATP molecules per glucose. Despite its high efficiency, OXPHOS is relatively slow and inherently generates reactive oxygen species (ROS) as byproducts, which can trigger oxidative damage to cellular components and promote apoptosis.

In contrast, glycolysis is a faster, less efficient process, yielding only 2 ATP per glucose, but produces lower levels of ROS, supporting cell survival under stress^1,2^. In addition, its end-product, pyruvate scavenges mitochondrial ROS, offering further anti-apoptotic benefits^3^. The metabolic reprogramming from OXPHOS to glycolysis, known as the Warburg effect, is a hallmark of cancer^4–7^ and is increasingly implicated in therapeutic resistance^3,4,8–10^, including resistance to EGFR tyrosine kinase inhibitors (TKIs).

When OXPHOS is compromised, cancer cells exhibit remarkable metabolic plasticity, reprogramming energy pathways to sustain bioenergetics and redox balance under drug pressure.

Notably, Ali et Levantini et al. recently identified a mutant-EGFR/FASN axis specific to TKI-resistant cells, wherein palmitoylated nuclear EGFR contributes to acquired resistance. This finding underscores the complexity of adaptive responses in osimertinib-resistant (OsiR) cells, involving coordinated genetic, transcriptional, and metabolic rewiring that extends beyond canonical resistance mutations.

Notably, Ali et Levantini et al. recently identified a novel mutant-EGFR/FASN axis specific to gefitinib-resistant cells, in which palmitoylated nuclear EGFR contributes to acquired resistance^11^. This finding reinforces the notion that resistant cells exhibit a complex adaptive interplay of genetic, transcriptional, and metabolic strategies collectively supporting their survival and proliferation, that extends well beyond canonical resistance mutations ^11–14^.

To capture these adaptations, integrative transcriptomic and untargeted metabolomic approaches have been applied across a wide range of cancer types, including lung^9,15,16^, pancreatic^17,18^, glioblastoma^19^, melanoma^20^, mesothelioma^21^, hepatocellular^22^, prostate ^23,24^, ovarian^25^, renal^26^, thyroid^27^, gastric ^28^, colorectal^29,30^ and breast cancers^31–34^, as well as hematological malignancies ^35–38^. These strategies, including analysis of the National Cancer Institute (NCI)-60 cell line panels ^39^ and other cancer cell line models^40,41^ have highlighted key metabolic pathways associated with drug resistance.

However, untargeted analyses may overlook unconventional or poorly annotated metabolites that fall outside standard annotations. Thus, integrating untargeted analyses with systematic, targeted quantification of specific metabolites, complemented by transcriptional and enzymatic profiling offers a more comprehensive view of tumor metabolism. This thorough strategy can potentially uncover critical blind spots and reveal alternative metabolic nodes that drive drug resistance, which may otherwise remain hidden in conventional bioinformatic-driven analyses.

One such underexploited yet highly informative strategy is exometabolomic profiling, the targeted analysis of metabolites in the extracellular medium (ECM). As a dynamic interface between cells and their microenvironment, the ECM reflects shifts in metabolic flux and offers real-time readouts of anabolic and catabolic activities ^42^. This methodology can therefore efficiently identify biomarkers linked to metabolic adaptations^42–44^. Importantly, this minimally invasive method preserves analyte integrity by avoiding cell lysis and matrix effects caused by extraction artifacts ^45–48^. Prior studies have employed cell culture *exometabolome* ^43,45–54^ to explore cancer metabolism ^55,56^ and intercellular communications favoring cancer formation and progression ^55,57^.

Here, we employed high-resolution chromatographic platforms, including reversed-phase liquid chromatography (HPLC-DAD and LC-HRMS) and GC-MS, to systematically quantify ECM metabolites ^45,58^ in parental (Par) and OsiR H1975 NSCLC cells, harboring the common L858R/T790M EGFR mutations. Our targeted analyses revealed a remarkable accumulation of acetate, along with elevated pyruvate, acetaldehyde and lactate in resistant cells. Pathway enrichment analysis of the exometabolome revealed upregulation of glycolysis/gluconeogenesis, pyruvate metabolism, nicotinate and nicotinamide metabolism, and several amino acid pathways. By integrating these findings with transcriptomic analysis and protein expression validation by Western blot, we uncovered a coordinated metabolic adaptation in OsiR cells that transcends genetic mutational events.^59^ We discovered a broader cellular strategy to preserve viability under therapeutic pressure through the reprogramming of energy and redox metabolism. Specifically, we observed a decoupling of glycolysis from downstream oxidative metabolism: although glycolytic activity is enhanced, as evidenced by the accumulation of pyruvate, lactate, acetaldehyde, and acetate, the subsequent utilization of these glycolytic products via the TCA cycle and PPP is attenuated, as indicated by the downregulation of key genes in these pathways. This suggests that glycolytic intermediates are diverted away from oxidative metabolism toward alternative fates that sustain energy (ATP production) and redox balance. Notably, acetate emerges as a major byproduct. delineating a distinct and novel pyruvate–acetaldehyde–acetate (PAA) axis, which appears critical to support redox homeostasis via NADPH production.

Recent studies suggest a *de novo* pathway for acetate production from pyruvate^60–70^. Liu et al. demonstrated in several model cell lines that acetate can be chemically generated through a non-enzymatic reaction, in which ROS directly attack pyruvate via nucleophilic attack, converting it into acetate^60^. In this reaction, pyruvate acts as a key ROS scavenger, a protective mechanism triggered under ETC dysfunction, which also preserves NADH for its essential role in reducing pyruvate to lactate^71^.

However, more than 90% of cellular acetate production appears to derive from neomorphic enzyme activity of keto acid dehydrogenases (KDHs)^60^, including pyruvate dehydrogenase E1 (PDHE1). These enzymes, located in both the inner mitochondrial membrane and the nucleus, depend on multiple cofactors, including pyruvate, CoA, NAD^+^, ROS, and thiamine/thiamine pyrophosphate (TPP)^60,72^. PDHE1 plays a key role by catalyzing the decarboxylation of pyruvic acid in a TPP- and magnesium-dependent reaction. Liu et al. proposed a model in which imbalances in substrates and cofactor concentrations can alter PDH function^60^, resulting in the production of pyruvate-derived acetaldehyde and acetate. Importantly, in metabolically hyperactive tumors, acetate is often released into the extracellular space^7^, supporting metabolic symbiosis among neighboring cancer cells^61^.

Under anaerobic conditions, PDH can also participate in fermentation processes. In organisms such as *Saccharomyces cerevisiae,* PDH contributes to ethanol production; in some vertebrates (e.g., goldfish and carp), similar fermentations under hypoxia, lead to ethanol generation ^73^.

Along with these changes in the concentration of metabolites strictly related to glycolysis and Warburg effect, we observed elevated levels of adenine, nicotinic acid, and multiple amino acids in OsiR cells, consistent with a major metabolic shift toward biosynthetic and anabolic metabolism.

In summary, our findings reveal a distinct reorganization of central carbon metabolism in osimertinib-resistant NSCLC cells, characterized by increased glycolysis, decreased oxPPP, and activation of the alternative PAA axis. This metabolic fingerprint not only defines the resistant phenotype but also unveils a metabolic vulnerability within the acetate-NADPH axis, offering new therapeutic opportunities against refractory disease.

## 2. Experimental

### 2.1. Chemicals and procedures

Sulfuric acid for HPLC analysis (30743, Honeywell Fluka, 95-97%) and methanol for RP-HPLC (34860, Merck, 99.9%) were used. Standard solutions for HPLC (TraceCERT®, 1000 mg/L in water) were purchased from Sigma-Aldrich (Milan, Italy). Analytes stock solutions were prepared by dissolving weighed amounts of pure compounds in deionized water and stored at 4°C for up to one month. All sample and solution preparations/dilutions were performed gravimetrically using ultrapure water (MilliQ; 18.2 MΟ cm-1 at 25 °C, Millipore, Bedford, MA, USA).

### 2.2. Cell cultures

H1975 parental (Par) and Osimertinib-resistant (OsiR) cell lines were cultured for 72 h in RPMI 1640 medium supplemented with 10% fetal bovine serum (FBS, Sigma-Aldrich) at 37 °C in a humidified incubator at 5% CO_2_. Cells were authenticated via DNA fingerprinting^74^ and tested negative for mycoplasma contamination.

### 2.3. Exometabolome analysis

Following 72 h incubation, the extracellular medium was collected from cultured cells and subjected to downstream analysis.

### 2.4. Pyruvate quantification by LC-HRMS

Pyruvate levels were quantified by LCMS. The analysis was carried out on an ultra-high-performance liquid chromatography (UHPLC; Vanquish Flex Binary pump) coupled to a diode array detector (DAD) and a high-resolution (HR) Q Exactive Plus mass spectrometer (MS), based on Orbitrap technology, equipped with a heated electrospray ionization (HESI) source (Thermo Fischer Scientific Inc., Bremen, Germany).

Chromatographic separation was achieved on a Zorbax® Eclipse Plus Phenyl-Hexyl column (Agilent, CA, United States; 4.6 x 250 mm, 5 µm). The flow rate was 0.8 mL/min with a 1:1 splitting system to the MS and DAD/UV detector. The column temperature was maintained at 40°C, and the injection volume was 2 μL.

Eluents consisted of MeOH/HCOOH 0.5% v/v (solvent B); H_2_O/HCOOH 0.5% v/v (solvent A) (all solvents were of ultra-purity grade, ROMIL Ltd, Cambridge, GB).

The gradient was set as follows: isocratic 100% A until 15 min, linear gradient from 0 to 80% (B) in the range 15-25 min, from 80% to 100% (B) in the range 25-35 min and isocratic 100%B for 2 min. The chromatographic run was complete in 44 min including re-equilibration in 100% solvent A. Ionization parameters were: nebulization voltage 3400 V (+) and 3200 V (−), capillary temperature 290°C, sheath gas (N_2_) 24 (+) and 28 (−) arbitrary units, auxiliary gas (N_2_) 5 (+) and 4 (−) arbitrary units, S-lens RF level 50. Scan mode: Full scan, Scan range: m/z 54-800. Resolution 70,000 at m/z 200. Polarity: positive and negative ionization mode.

Data were acquired and analyzed by Xcalibur 3.1 software (Thermo Fischer Scientific Inc., Bremen, Germany). LCMS files were processed using Xcalibur 4.1 (Thermo Fisher Scientific). Extracted ion chromatograms (EICs) were obtained by exact mass range calculations based on the elemental formula (C_n_H_n_O_n_P_a_N_n_S_n_ - H for negative ions +H for positive ions), with ± 3 ppm (ΔM/M) *10^6^ of accuracy. Peaks were manually integrated, and results elaborated in Microsoft Excel.

### 2.5. Lactate quantification by RP-HPLC-DAD

Lactate concentrations in the extracellular medium were measured by reversed-phase liquid chromatography (RP-HPLC) equipped with diode array and fluorescence detectors (HPLC-DAD)^74^. Lactate was quantified by LC-DAD to avoid saturation of the MS detector, given its high concentration (in the mM range) in the extracellular compartment. For RP-HPLC-DAD analysis, samples were diluted 1:5 in 5 mM sulfuric acid, filtered through 0.20 mm RC Mini-Uniprep filters (Agilent Technologies, Italy), and injected (V_inj_ = 5 μL) into the HPLC system. The same chromatographic column used for LC-HRMS analysis was employed. Lactate was identified by comparison with the retention time and UV spectra of standard compounds. The signals were manually integrated, and concentrations were determined using a calibration curve generated with corresponding analytical standards.

### 2.6. Acetate and acetaldehyde quantification by HS-GC-MS

Acetate and acetaldehyde levels were quantified by headspace (HS) GC-MS. Acetaldehyde was determined as such. Acetate was determined after acidification with H_2_SO_4_ to convert it to volatile acetic acid via (HS) GC-MS.

For acetate analysis, 500 μL of sample or standard acetate (0, 50, 100, 250, 400 μg/mL; prepared gravimetrically from Certified Standard), 50 μL of ^2^H_3_ acetate at 1000 μg/mL (Internal Standard), and 100 μL of 5 M H_2_SO_4_ were added to 10 mL HS vials and sealed immediately. Standards and samples were prepared in triplicate. Calibration curves were prepared with area at m/z 43 /area at m/z 46 ratio plotted against [acetate]/[acetate ^2^H_3_] concentration ratio, using Agilent MassHunter Quantitative Analysis Ver. 10.2.

A GC Agilent 6850 together with an Agilent single quadrupole MS 5975c, equipped with an Agilent GC 80 CTC PAL ALS, was employed for the measurements. For acetate the separations were achieved using a J&W DB-WAX-UI capillary column (30.0 m, 250.00 μm internal diameter, 0.50 μm film thickness) with a constant flow of 1.0 mL/min (average velocity: 36 cm/sec; incubation at 80 °C for 600 s; 1000 μL injection volume). The oven program was as follows: initial temperature 50 °C, hold 5 min, 10.00 ° C/min up to 150 °C, hold 2.00 min, 20 °C/min up to 240 °C, hold 8.5 min (total time 30 min). Pulsed Splitless inlet mode; initial temperature 200 °C, pressure 45.4 kPa, pulse pressure 120 kPa, pulse time 0.10 min, purge flow 200.0 mL/min, purge time 0.10 min, total flow 203.7 mL/min, gas type helium; transfer line temperature 250 °C; SIM acquisition mode. MS conditions were as follows: EI ionization mode at 70eV electron energy; Ion Source Temp. 250 °C Quadrupole Temp. 150 °C; Acquisition mode SIM; acquisition m/z 43, 46, 60, 63; Dwell Time 100 ms; quantification ions: 43 and 46; qualifier ions: 60 and 63. Acquisition Software: Agilent MSD ChemStation Ver. E.02.02.

For acetaldehyde analysis, 500 μL of sample or standard acetaldehyde (0, 50, 100, 250, 400 μg/mL; prepared gravimetrically from Certified Standard) were spiked with 50 μL of ^2^H_3_ ethanol at 89.2 μg/mL (Internal Standard). For acetaldehyde the separations were achieved using the same column with a constant flow of 1.0 mL/min (average velocity: 36 cm/sec; incubation at 25 °C for 600 s; 1000 μL injection volume). The oven program was as follows: initial temperature 25 °C, hold 5 min, 25.00 °C/min up to 240 °C, hold 5.00 min. Quantification ions: 29.44 for acetaldehyde, 45 and 46 for ethanol, and 48, 49 for ^2^H_3_. The detailed description has been previously reported^75^.

### 2.7. SDS-PAGE and Western Blotting

For protein analysis, 15 μg of protein from each sample (Par and OsiR cells) was separated by SDS-PAGE on 10% gel and subsequently transferred to nitrocellulose membranes with the TransBlot Transfer System (Bio-Rad). Membranes were blocked for 1 hour at room temperature in either 5% non-fat dry milk or 5% bovine serum albumin (BSA) diluted in Tris-buffered saline with 0.1% Tween-20 (1X TBST). Primary antibodies (listed below) were incubated overnight at 4°C. After incubation, membranes were washed three times for 10 minutes each with 1X TBS-T at room temperature. HRP-conjugated secondary antibodies (listed below) were then applied for 1 hour at room temperature. Proteins bands were detected by chemiluminescence (Pierce ECL Western Blotting Substrate Cat #32106) and visualized using the Chemidoc Imaging System (Biorad). Membranes were subsequently stripped using Stripping Buffer Solution (Himedia, #ML163), as per the manufacturer’s instructions, and re-probed with an anti-β-actin mouse antibody to verify equal protein loading. After three washes with 1X TBS-T, membranes were incubated with the appropriate anti-mouse secondary antibody (details below).

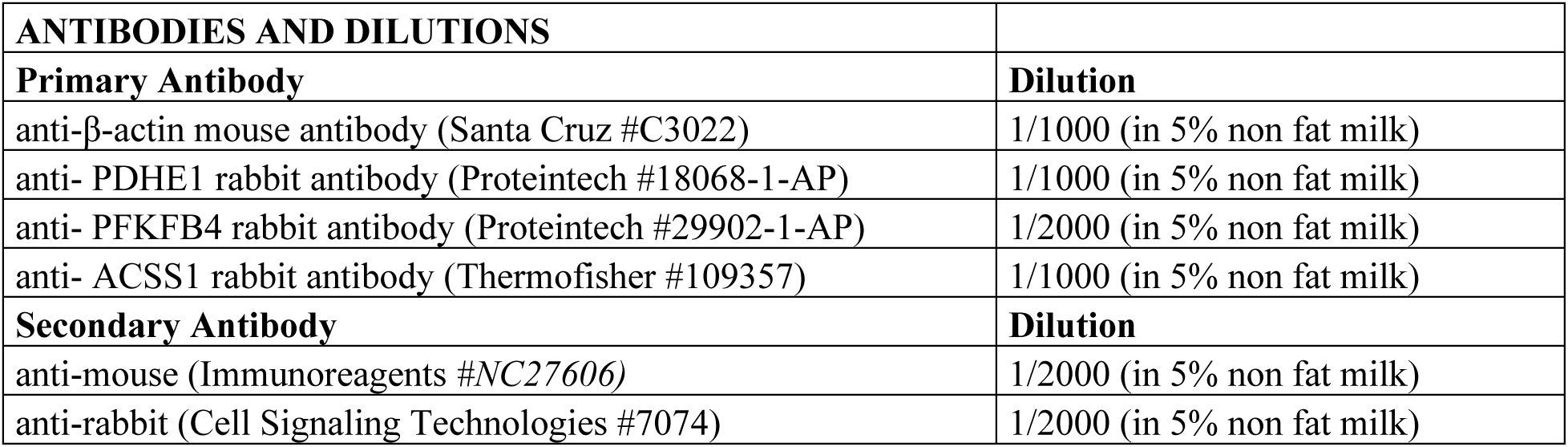

### 2.8. Pathway analysis

Pathway topology and biomarker analysis of discriminating metabolites was carried out by using MetaboAnalyst 6.0.^76,77^

## 3. Results

### 3.1 Exometabolomic Profiling by LC-HRMS, LC-DAD and GC-MS: Metabolic Indicators in Osimertinib-resistant cells

**Figure 1(A)** shows the results of pathway analysis of significantly altered exometabolites in OsiR *vs* Par cells determined using LC-HRMS, LC-DAD and GC-MS. Pathways are mapped via the KEGG database and displayed according to pathway impact and adjusted *p*-value (expressed as − log(*p* value)), highlighting the most altered metabolic pathways distinguishing OsiR from Par cells. Analogous results were obtained through metabolite set enrichment analysis (MSEA) (**Figure 1B**), a metabolomics approach that evaluates the enrichment of predefined metabolic pathways to identify those most significantly perturbed ^76,77^.

**Figure 1.**
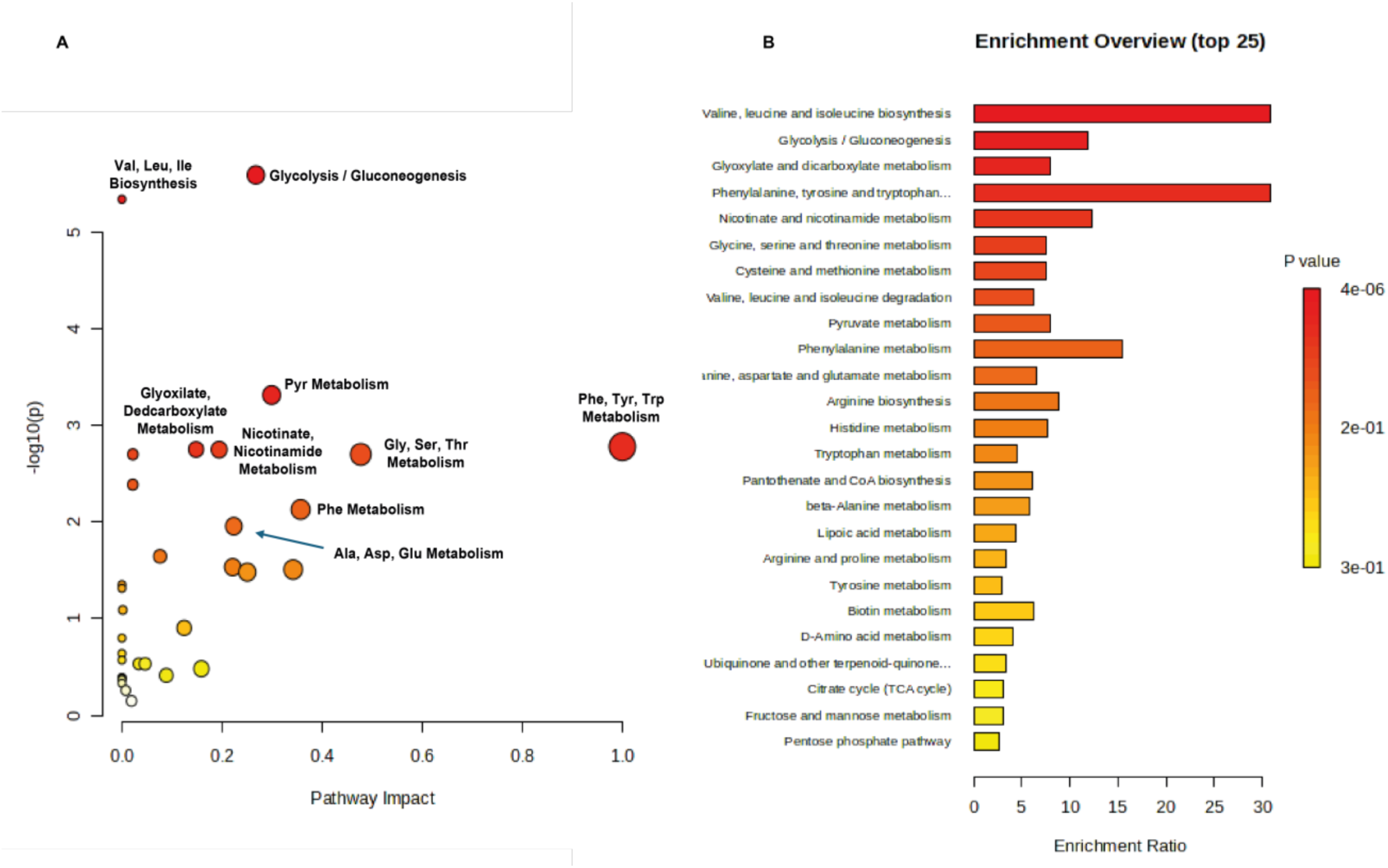
**(A)** Pathway analysis of significantly altered exometabolites in OsiR *vs* Par cells using LC-HRMS, LC-DAD and GC-MS. Pathways are mapped via the KEGG database and displayed according to pathway impact and adjusted *p-*value. Dot size = pathway impact; color = statistical significance (darker = more significant). **(B)** Metabolite set enrichment analysis (MSEA) confirming the most impacted pathways in OsiR vs Par cells. Statistical and topological analyses were performed using MetaboAnalyst 6.0^76,77^. Pathway impact represents a combination of pathway enrichment and topology analysis (centrality). Circle indicates the pathway impact, while color intensity represents the statistical significance (more intense red color corresponds to lower *p*-values, and thus higher significance).

The pathway topology and biomarker analysis identified several significantly dysregulated pathways in OsiR cells, including glycolysis/gluconeogenesis, pyruvate metabolism, phenylalanine, tyrosine and tryptophan biosynthesis, glyoxylate and dicarboxylate metabolism, nicotinate and nicotinamide metabolism, and phenylalanine metabolism (**Table 1**).

**Table 1.** Top 10 significantly dysregulated metabolic pathways in OsiR cells.

| | Total | Expected | Hits | Raw $p$ | $-\log_{10}(p)$ | Holm adjust | FDR | Impact |
| --- | --- | --- | --- | --- | --- | --- | --- | --- |
| Glycolysis / Gluconeogenesis | 26 | 0.44571 | 6 | 2.5748E-06 | 5.5893 | 0.00020598 | 0.00018353 | 0.2677 |
| Valine, leucine and isoleucine biosynthesis | 8 | 0.13714 | 4 | 4.5883E-06 | 5.3383 | 0.00036248 | 0.00018353 | 0 |
| Pyruvate metabolism | 23 | 0.39429 | 4 | 0.00048638 | 3.313 | 0.037937 | 0.01297 | 0.29918 |
| Phenylalanine, tyrosine and tryptophan biosynthesis | 4 | 0.068571 | 2 | 0.0016632 | 2.779 | 0.12807 | 0.019978 | 1 |
| Glyoxylate and dicarboxylate metabolism | 32 | 0.54857 | 4 | 0.0017764 | 2.7505 | 0.13501 | 0.019978 | 0.14815 |
| Nicotinate and nicotinamide metabolism | 15 | 0.25714 | 3 | 0.0017838 | 2.7487 | 0.13501 | 0.019978 | 0.1943 |
| Cysteine and methionine metabolism | 33 | 0.56571 | 4 | 0.0019978 | 2.6995 | 0.14783 | 0.019978 | 0.02184 |
| Glycine, serine and threonine metabolism | 33 | 0.56571 | 4 | 0.0019978 | 2.6995 | 0.14783 | 0.019978 | 0.47737 |
| Valine, leucine and isoleucine degradation | 40 | 0.68571 | 4 | 0.0041085 | 2.3863 | 0.29581 | 0.03652 | 0.02168 |
| Phenylalanine metabolism | 8 | 0.13714 | 2 | 0.007439 | 2.1285 | 0.52817 | 0.059512 | 0.35714 |

Specific metabolites related to glycolytic pathway were quantified in the exometabolome of Par and OsiR cell lines cultured for 72 h. **Table 2** summarizes the nM concentrations (mean ± SD) of metabolites (n=3 biological replicates) measured in the ECM of Par and OsiR cell cultures, normalized to cell numbers.

**Table 2.**
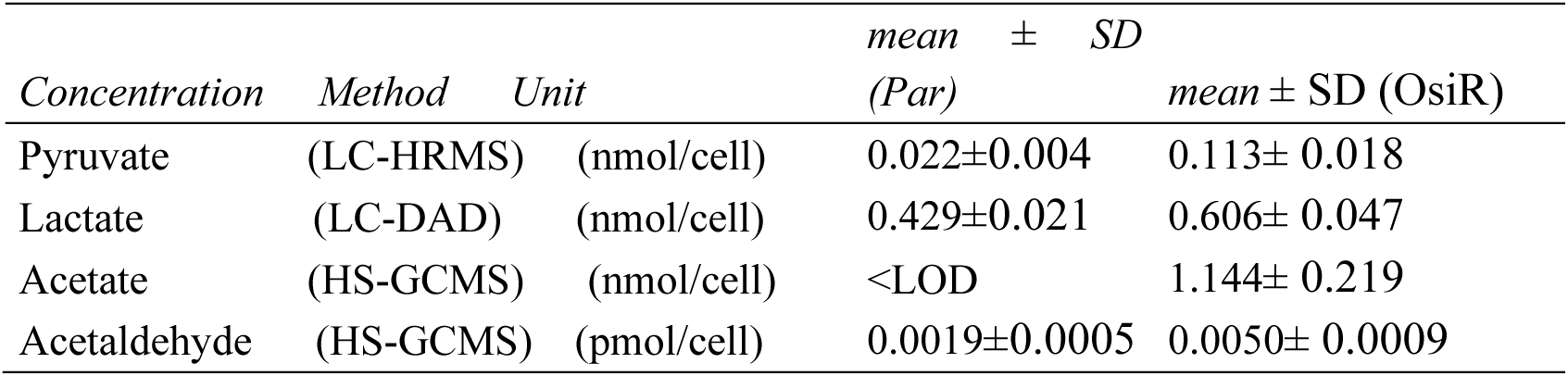
Concentration levels of pyruvate, lactate, acetate and acetaldehyde in the ECM of Par and OsiR cell cultures. Measurements were performed on three biological replicates. Values are normalized to cell numbers. LOD= limit of detection.

| Concentration | Method | Unit | mean ± SD<br>(Par) | mean ± SD (OsiR) |
| --- | --- | --- | --- | --- |
| Pyruvate | (LC-HRMS) | (nmol/cell) | 0.022±0.004 | 0.113± 0.018 |
| Lactate | (LC-DAD) | (nmol/cell) | 0.429±0.021 | 0.606± 0.047 |
| Acetate | (HS-GCMS) | (nmol/cell) | <LOD | 1.144± 0.219 |
| Acetaldehyde | (HS-GCMS) | (pmol/cell) | 0.0019±0.0005 | 0.0050± 0.0009 |

Untargeted metabolomics analysis via LC-HRMS further revealed significant differences in the levels of additional low-MW metabolites in the ECM of Par and OsiR cells.

**Figure 2** shows LC-HRMS peak areas/concentrations of metabolites related to the glycolysis/gluconeogenesis, significantly different in the ECM of Par and OsiR cells.

**Figure 2.**
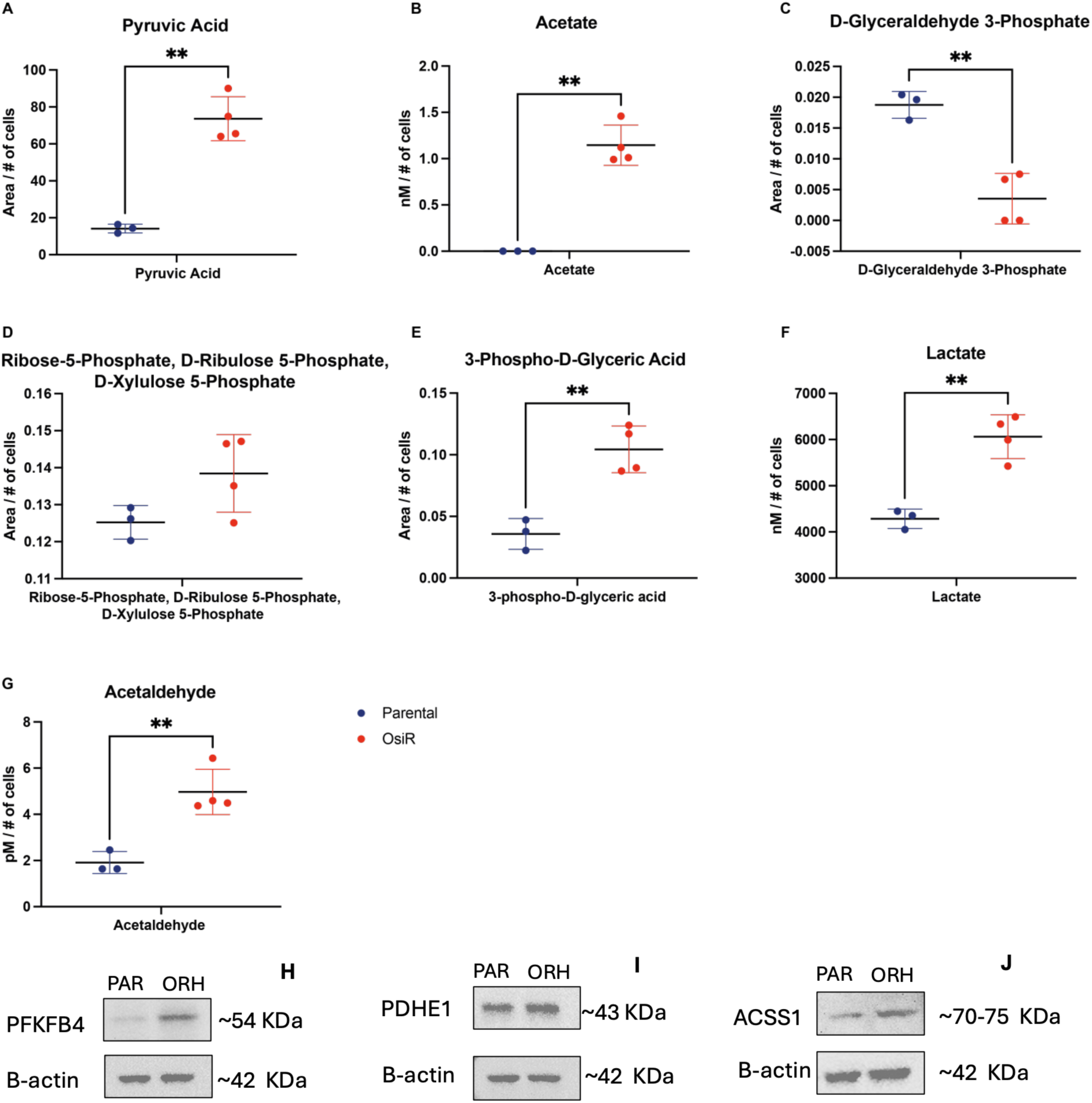
(A)-(G) Bar plot showing the peak areas/concentrations of pyruvate, acetate, lactate, acetaldehyde, and other metabolites associated with the glycolysis/gluconeogenesis, identified by chromatographic techniques (LC-HRMS, HPLC-DAD and HS-GCMS) in Par (blue dots) and OsiR ECM (red dots). Statistical significance: *p*< 0.001 (**); *p*<0.05 (*). (H)-(J) Western blot analyses of the human H1975 cell line (Par and OsiR). Protein lysates were immunoblotted with an anti-PFKFB4, anti-PDHE1 and anti-ACSS1. Loading was assessed with an anti-β-actin antibody. Expected size is shown in kDa.

To validate the metabolic reprogramming suggested by the exometabolomic analysis, the protein expression of key enzymes involved in glycolysis, PPP, the TCA cycle, and aldehyde detoxification was assessed by Western blot analysis, as recently reported,^78^ and as reported in **Figure 2 panels (H)-(J)**. OsiR cells exhibited elevated levels of transketolase (TKT), which bridges glycolysis and the PPP, TCA enzymes (pyruvate dehydrogenase complex, PDH, pyruvate dehydrogenase E1, PDHE1, specifically involved in the decarboxylation of pyruvate, and succinate dehydrogenase, SDH), enzymes involved in aldehyde detoxification Aldehyde Dehydrogenase 2 (ALDH2), and Aldehyde Dehydrogenase 7 Family Member A1 (ALDH7A1), and the glycolytic enzyme (aldolase A, ALDOA and 6-Phosphofructo-2-Kinase/Fructose-2,6-Biphosphatase 4, PFKFB4). Western blot analysis evidenced also increased levels of Acyl-CoA Synthetase Short-chain family member 1 (**ACSS1, Figure 2J**), catalyzing the conversion of acetate to acetyl CoA.

No significant differences in the expression of nicotinamide nucleotide transhydrogenase (NNT) (data not shown) were observed between Par and OR cells. NNT is an integral protein of the inner mitochondrial membrane catalyzing the coupling of the hydride transfer between NADH and NADP^+^ to proton translocation across the inner mitochondrial membrane. This enzyme uses energy from the mitochondrial proton gradient to produce high concentrations of NADPH, a critical reducing equivalent.

**Figure 3** shows LC-HRMS peak areas/concentrations of amino acids, metabolites related to amino acid metabolism, and NAD^+^/NADP^+^ biosynthesis significantly different in the ECM of Par and OsiR cells. The statistical significance was determined at *p*< 0.001 (**); and *p*<0.05 (*).

**Figure 3.**
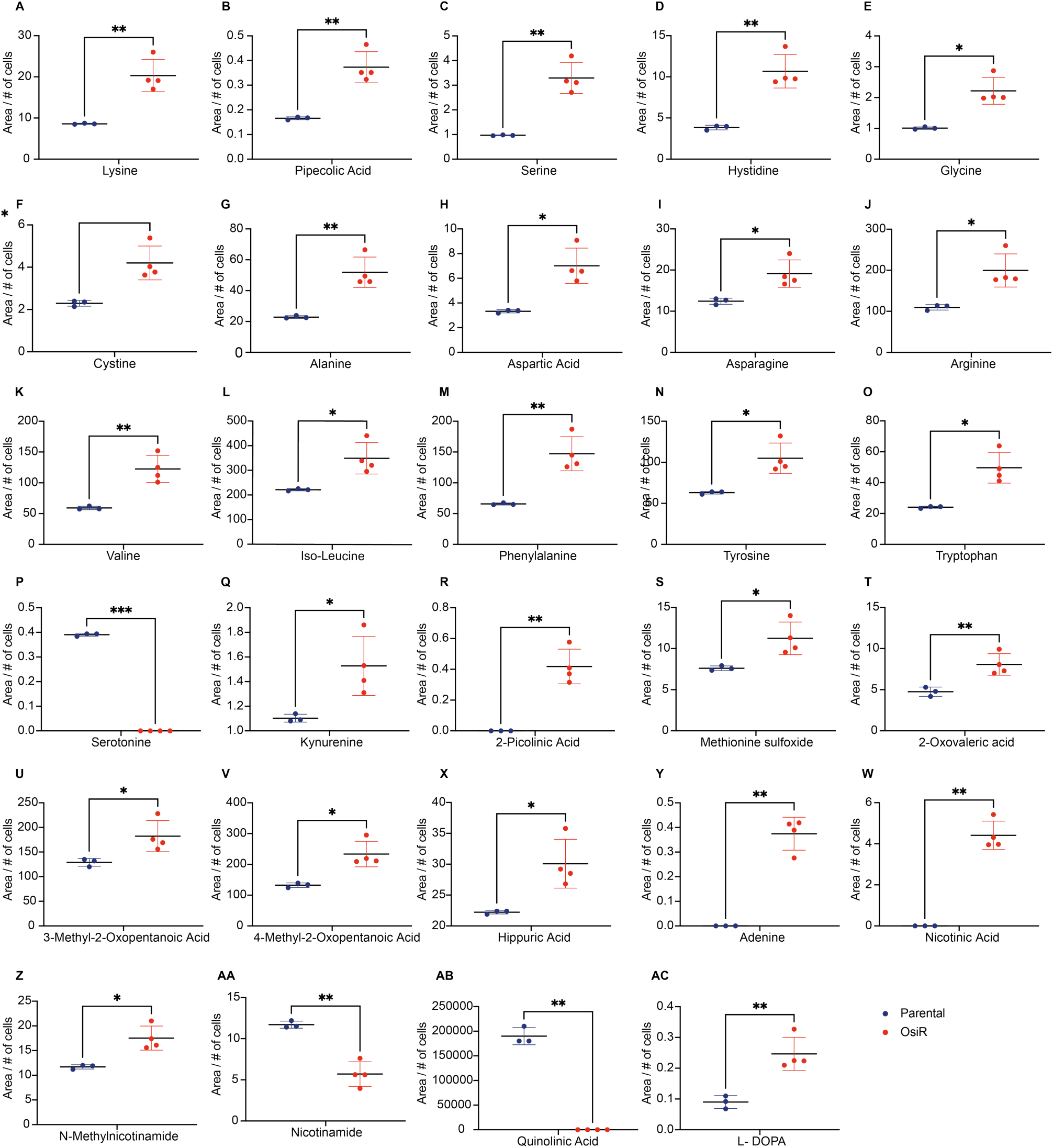
Bar plot showing the LC-HRMS peak areas of amino acids, metabolic intermediates, and NAD^+^/NADP^+^ biosynthesis in Par (blue dots) and OsiR ECM (red dots). Statistical significance: *p*< 0.001 (**); *p*<0.05 (*).

## 4. Discussion

Resistance to osimertinib in EGFR-mutant NSCLC poses a significant challenge. This study examines the metabolic reprogramming associated with this resistance by integrating exometabolomic profiling of parental H1975 cells and their resistant counterparts with transcriptomic and targeted protein analysis.

Bioinformatic analysis of our metabolomic results revealed substantial metabolic reprogramming in OsiR cells, identifying, as expected, alterations in core bioenergetic pathways, such as glycolysis/gluconeogenesis, pyruvate metabolism, glyoxylate and dicarboxylate metabolism, as well as vitamin and amino acid metabolism (**Figure 1** and **Table 1**). Transcriptomic data and complementary biological assays consistently highlighted the involvement of the aforementioned metabolic pathways. Altered mitochondrial function and morphology, impaired ETC activity, and compromised OXPHOS, likely due to mtDNA mutations or changes in gene expression, have been reported^78^. Collectively, these alterations contribute to the metabolic reprogramming observed in OsiR cells^78^.

### Glycolysis, pentose phosphate pathway and tricarboxylic acid switch

The interplay between glycolysis, the PPP and the TCA cycle provides cancer cells with remarkable metabolic flexibility, enabling them to dynamically switch between these pathways as needed, by rerouting carbon flux where needed. Among these pathways, the PPP, which branches from glycolysis, plays a dual function: generating NADPH for redox homeostasis and reductive biosynthesis, and producing ribose-5-phosphate for nucleotide, vitamin and cofactor biosynthesis. The oxidative phase of the PPP (ox-PPP), initiated by glucose-6-phosphate dehydrogenase (G6PDH), converts glucose-6-phosphate (G6P) into ribulose-5-phosphate while generating NADPH. This branch is particularly active in cells requiring high levels of NADPH. Following the ox-PPP, the non-oxidative phase of the PPP (non-ox-PPP), driven by TKT and TALDO, interconverts sugar phosphates to support nucleotides and amino acids biosynthesis. The flexibility of the PPP enables cells to adapt to varying metabolic needs while maintaining a balance between NADPH production and ribose-5-phosphate biosynthesis. These pathways are tightly interconnected with glycolysis and gluconeogenesis, enabling dynamic adaptation to metabolic stress. Metabolic intermediates from the PPP can reintegrate into glycolysis or feed into gluconeogenesis, highlighting the integrated and adaptable nature of cellular carbon metabolism in cancer.

Pyruvate, mainly derived from glycolysis, serves as a key metabolic node at the heart of this network, linking glycolysis to the TCA cycle. Pyruvate also derives from additional sources including the conversion of malate (via malic enzymes), alanine (via alanine transaminase, ALT), and other gluconeogenic amino acids ^79^. Once formed, pyruvate is a critical metabolic junction. Pyruvate can either remain in the cytosol and be reduced to lactate by lactate dehydrogenase (LDH), typical of the Warburg effect, or be transported into the mitochondrial matrix to fuel the TCA cycle as acetyl-CoA through the action of the pyruvate dehydrogenase complex (PDH).

In cancer cells, uncontrolled proliferation and resistance to therapy are commonly linked to the Warburg effect, characterized by increased glycolysis and lactate production even in the presence of oxygen ^3,8,10,80–83^. In OsiR cells, elevated levels of pyruvate (**Figure 2A**) align with this metabolic shift, indicating an increase glycolytic flux that redirects pyruvate towards lactate production, consistent with the Warburg profile. However, the metabolic phenotype of OsiR cells is not limited to this classical pathway and appears to extend beyond the Warburg effect. Emerging evidence points to additional metabolic adaptations beyond classical glycolysis. Notably, the accumulation of acetate (**Figure 2B**) in the extracellular milieu of OsiR cells, suggests that a portion of pyruvate is diverted away from lactate and processed through alternative metabolic routes under osimertinib-induced stress. In this context, acetate may behave as a byproduct but also as a carbon source for nearby cancer cells, reinforcing metabolic symbiosis within the tumor environment.

### Hexokinases and Glycolytic Commitment

Among the key regulatory enzymes, **hexokinases (HK)** are high-affinity enzymes physically associated with the outer mitochondrial membrane, which catalyze the first and irreversible step of glycolysis by phosphorylating glucose to glucose-6-phosphate (G6P). Reflecting metabolic adaptation, their expression varies with resistance, as **HK1 is upregulated** (+0.424-fold, adj. *p*-value 2.03 ×10^−7^), while **HK2 is downregulated** in OsiR cells (−0.299-fold, adj. *p*-value 3.19 ×10^−5^). HK1, considered the housekeeping isoform^84,85^, ensures glucose is efficiently trapped within the cell and committed to the glycolytic pathway, providing a steady supply of G6P for both glycolysis and the PPP. In contrast, HK2, a HIF1 transcriptional target, commonly contributes to metabolic rewiring under hypoxia^84^.

Once phosphorylated, glucose enters either glycolysis or the PPP. In OsiR cells, both hexose-6-phosphate dehydrogenase (**H6PDH)**, localized in the microsomes^86^, and glucose-6-phosphate dehydrogenase **(G6PDH),** localized in the cytosol, are downregulated (−0.813-fold, adj. *p*-value 9.34 ×10^−8^ and −0.579-fold, adj. *p*-value 3.35 ×10^−4^, respectively). These enzymes catalyze the first committed step of the ox-PPP. Their downregulation is accompanied by decreased expression of 6-phosphogluconolactonase (**PGLS**) (−0.546-fold, adj. *p*-value 1.01 ×10^−4^) and 6-phosphogluconate dehydrogenase (**PGD**) (+0.157-fold, adj. *p*-value 1.16 ×10^−3^), the latter being the second dehydrogenase in the PPP. As a result, the activity of the ox-PPP is reduced in OsiR cells. This metabolic reprogramming leads to a preferential routing of glucose through glycolysis, reflecting the Warburg effect, rather than its redirection to the ox-PPP. This adaptation eventually provides the cell with rapid ATP generation through glycolysis, which yields less ATP per molecule of glucose compared to OXPHOS, but it does so at a faster rate, allowing for rapid energy production. However, this shift compromises the production of NADPH, which is typically produced via the ox-PPP. Despite this, resistant cells maintain aggressive phenotypes, including their migratory and invasive capabilities, implying the presence of compensatory mechanisms that support their thriving under stress.

### oxPPP Enzymes and Drug Resistance

The role of G6PDH is crucial for cancer in mediating drug resistance, as it provides NADPH, an essential cofactor to buffer oxidative stress ^87^. Mutations in G6PDH, the rate-limiting enzyme of the PPP, decrease NADPH levels and increase ROS, diminishing metastatic potential ^88^. Conversely, PGD upregulation is seen in imatinib-resistant gastrointestinal stromal tumors (GISTs), where it promotes cell proliferation and inhibits apoptosis^89^. More broadly, the oxidative branch of the PPP is consistently implicated in cancer drug resistance, with overexpression and heightened activity of PPP enzymes contributing to drug resistance across multiple cancers ^90–92^.

### Phosphofructokinase Regulation

The first uniquely committed step of glycolysis, the conversion of fructose 6 phosphate (F6P) to fructose 1,6 biphosphate (F1,6BP), is catalyzed by **phosphofructokinase** (PFK1). This tetrameric enzyme exists in three isoforms: muscle (PFKM), liver (PFKL), and platelet (PFKP). The PFK1 reaction is irreversible and F1,6BP is only used for glycolysis. In OsiR cells, transcriptomic analysis revealed **upregulation of PFKM** (+0.257-fold, adj. *p*-value 2.70 ×10^−5^) and **downregulation of PFKL** (−0.326-fold, adj. *p*-value 8.68 ×10^−6^) suggesting the predominant PFK1 complex is exclusively composed by PFKM subunits. This configuration enhances glycolysis and increases F1,6BP levels. PFK1 is regulated by fructose-2,6-bisphosphate (F2,6BP), a potent activator of PFK1 that dictates its activity and the glycolytic rate^93,94^.

In OsiR cells, both **PFKFB4** and **PFKFB3**, isoenzymes of 6-phosphofructo-2-kinase/fructose-2,6-bisphosphatase are **upregulated** (+0.465-fold, adj. *p*-value 1.72 ×10^−2^ and +0.149-fold, adj. *p*-value 1.51 ×10^−2^, respectively), controlling the cytoplasmic levels of F2,6BP, which allosterically activates PFK1, thus eventually enhancing glycolytic flux^95^. Western Blot analysis confirmed the upregulation of PFKFB4 at the protein level (**Figure 2K**). PFKFB4, particularly increased in lung adenocarcinoma, promotes oncogenic phenotypes^96^ by redirecting glucose flux to the non-ox-PPP (via transketolase, TKT), facilitating nucleotide, amino acid, and vitamin biosynthesis^96^. PFKFB3, though more moderately upregulated, supports glycolysis by directing PFK1 activity This dual role helps cancer cells balance energy production, redox homeostasis, and biosynthetic needs under drug resistance conditions. PFKFB4’s higher fructose-2,6-bisphosphatase activity, compared to PFKFB3, helps to degrade F2,6BP, redirecting glucose flux to the non-ox-PPP ^97,98^. Overall, these metabolic adaptations, favoring the coupling of glycolysis with the non-ox-PPP, are relevant for cellular adaptation, providing cancer cells with metabolic flexibility needed to adapt and resist therapy.

Supporting this glycolytic upregulation, key glycolytic enzymes such as **aldolase A (ALDOA)** (+0.497-fold, adj. *p*-value 1.52 ×10^−2^) and **enolase isoforms (ENO1 and ENO2)** are also overexpressed in OsiR cells (+0.219-fold, adj. *p*-value 1.35 ×10^−5^ and +0.910-fold, adj. *p*-value 9.06 ×10^−6^, respectively). **ALDOA** catalyzes the reversible conversion of F1,6PB to glyceraldehyde-3-phosphate (G3P) and dihydroxyacetone phosphate (DHAP), a critical glycolytic node, while and **ENO1 and ENO2** catalyze the reversible conversion of 2-phosphoglycerate (**2PG**) to phosphoenolpyruvate (PEP).

ALDOA was found upregulated in chemoresistance of breast cancer, enhancing glycolysis and the non-oxidative branch of the PPP ^99^. The increased expression of ENO2 is also significant because of its non-glycolytic functions, including the modulation of epithelial-mesenchymal transition (EMT) and drug sensitivity, contributing to therapy resistance^100–102^.

LC-HRMS analysis evidenced the **decrease in G3P (Figure 2C) in OsiR cells.** G3P levels may decrease as it is shunted to the non-ox-PPP by TKT. TKT converts (i) G3P and F6P into xylulose-5-Par and erythrose-4-phosphate, and (ii) G3P and sedoheptulose-7-Par into D-xylulose-5-Par and ribose-5-Par. Although transcriptomics shows that **TKT** is equally expressed in Par and OsiR cells, Western blot analysis^78^ evidenced TKT overexpression in OsiR cells. These data align with the levels of ribose-5-phosphate, D-ribulose 5-phosphate, and D-Xylulose 5-phosphate that tends to increase in OsiR ECM (**Figure 2D**). Ribose-5-Par is mostly relevant in proliferative cells for the biosynthesis of purine and pyrimidine nucleotides, as well as for the active (coenzyme) forms of several vitamins, such as NAD, NADP, FAD, CoA, in any cell type.^103^

Thus, in OsiR cells, the enhanced glycolysis and TKT activity supports the non-ox-PPP and nucleotide biosynthesis^99^. In cancer cells, PPP activity is elevated ^104–108^, and the enzyme transketolase (TKT) is overexpressed in various cancers, including lung cancer. TKT is essential for the non-oxidative branch of the PPP and significantly impacts cancer progression and patient survival ^109–115^. Recent studies show that TKT is upregulated in NSCLC cells, and its knockdown inhibits cell proliferation and enhances the efficacy of gefitinib, highlighting its potential as a therapeutic target in advanced lung cancer treatment ^112^.

In OsiR cells, G3P is also converted to form **3-phosphoglycerate (3PG)** that has been observed by LC-HRMS to **increase in OsiR (Figure 2E)**. Phosphoglycerate mutase 1 **(PGAM1)** reversibly converts 3PG to 2PG, a crucial step in glycolysis. This step is unique because most glycolytic intermediates involved in anabolic biosynthesis lie upstream of it^116^. Interestingly, 3PG inhibits PGD in the ox-PPP, while 2-PG activates the 3-phosphoglycerate dehydrogenase (PHGDH), which catalyzes the conversion of 3PG into 3-phosphohydroxypyruvate, the committed step in L-serine biosynthesis. A recent in vitro and in vivo study revealed increased levels of 3PG in cancer cells and demonstrated the significant role of PHGDH^117^ in cancer progression.

Lactate dehydrogenase (LDH) catalyzes the conversion of L-lactate and NAD^+^ to pyruvate and NADH. Its isoenzymes A and B are known to be involved in the control of T cell glycolysis and differentiation^118^. In contrast, **LDHC** is abundant in spermatozoa, which utilize aerobic glycolysis to meet their energy requirements^119^. In OsiR cells, LDHC isoform is upregulated (+0.975-fold, adj. *p*-value 1.09 ×10^−5^) and preferentially expressed over LDHA (−0.2)^78^. LDHC’s high catalytic activity at acidic pH and elevated temperatures supports the Warburg effect^120^. The overexpression of LDHC matches with the high concentration of **lactate** in ECM found in OsiR cells (**Figure 2F** and **Table 2**). Lactate concentration in OsiR is 0.606±0.047 nmol/cell, which is significantly higher than lactate concentration in Par cells (0.429±0.021 nmol/cell) (**Table 2**).

Among **TCA-related genes,** Par and OsiR show significant differences in the expression of five genes: aconitase (ACO1), succinate dehydrogenase (SDH), succinyl-CoA ligase ADP-forming subunit beta (SUCLA2), succinate-CoA ligase GDP-Forming subunit beta (SUCLG2), isocitrate dehydrogenase (IDH), adenosine triphosphate citrate lyase (ACLY), as well as the pyruvate dehydrogenase complex (PDH).

In OsiR we observe also the upregulation of the gene of mitochondrial **aconitase (ACO1 also known as IRP1)** (+0.700-fold, adj. *p*-value 1.04 ×10^−7^) that is required for the TCA cycle catalyzing the citrate/isocitrate interconversion, but also independently required for the mitochondrial genome maintenance and other unknown functions, such as iron homeostasis^121^. When cellular iron levels are high, ACO1 binds to a 4Fe-4S cluster and functions as an aconitase.

Transcriptomics evidenced the upregulation of several gene encoding for SDH (**SDHC**, SDHB, and SDHAF1) (+0.688-fold, adj. *p*-value 1.53 ×10^−5^, +0.156-fold, adj. *p*-value 2.07 ×10^−2^, and +0.443-fold, adj. *p*-value 1.40 ×10^−2^, respectively) and Western blot confirmed the hyperexpression in OsiR of SDH (**Figure 4B**). SDH, or mitochondrial complex II (MCII), is composed of four subunits and is the only enzyme participating in both the TCA cycle and the ETC, catalyzing the oxidation of succinate and ubiquinone to fumarate and ubiquinol, while transferring electrons as part of the TCA cycle to OXPHOS. Interestingly, it does not show mutations in OsiR^78^. While the other complexes in the ETC contain some polypeptide subunits encoded by the mitochondrial genome, all four subunits of MCII are encoded by nuclear genes, thus making the respiratory chain defects in MCII relatively rare^122^. **SDHC** plays an important role in energy generation to maintain cellular growth^122^; its overexpression has been associated with increased ROS production and defects in SDH assembly^123^. **SDHB** encodes the iron-sulfur subunit of complex II(Ip)^124^, and **SDHAF1**, a mitochondrial assembly factor, is essential for SDH assembly, though it does not physically associate with the complex *in vivo*. The SDH complex also plays a role in oxygen-sensing mechanism by converting succinate, an oxygen-sensor metabolite that stabilizes hypoxia-inducible factor 1 (HIF1)^125^.

Interestingly, two genes encoding succinyl-CoA synthetase show transcriptional changes: **SUCLG2** (+0.236-fold, adj. *p*-value 7.28 ×10^−4^) is upregulated, while SUCLA2 (−0.291-fold, adj. *p*-value 1.18 ×10^−4^) is downregulated in OsiR cells. SUCLG2 encodes the GTP-specific beta subunit of succinyl-CoA synthetase, and is implicated in mitochondrial dysfunction and succinylation regulation in lung adenocarcinoma^126^.

**SUCLA2** encodes an ATP-specific SCS beta subunit that dimerizes with the succinyl-CoA synthetase alpha subunit. However, it is reported that the metabolic contribution of SUCLA2 during metastasis is independent of its role in the TCA cycle and it is a critical factor that manages redox balance and promotes survival of disseminated cancer cells during metastasis^127^.

**Isocitrate dehydrogenase** (IDHs) catalyzes the oxidative decarboxylation of isocitrate to α-KG. These enzymes belong to two distinct subclasses, one of which utilizes NADP^+^ as the electron acceptor. IDH-encoding genes are downregulated in OsiR cells, including **IDH1** (−0.389-fold, adj. *p*-value 8.69 ×10^−6^) located in the cytoplasm and peroxisome, and mitochondrial **IDH2** (−0.925-fold, adj. *p*-value 4.93 ×10^−6^). IDH plays a role in intermediary metabolism and energy production and may tightly associate or interact with the PDH complex. In OsiR, the NAD^+^-dependent non-catalytic subunit gamma of IDH, **IDH3G** (−0.631-fold, adj. *p*-value 4.91 ×10^−7^), which can be allosterically activated by citrate or/and ADP, is also downregulated, suggesting the redirection of **citrate** toward the cytoplasm, where it can be the substrate of **ACLY. SLC25A1**, the gene that codes for a protein that transports citrate across the mitochondrial membrane also known as mitochondrial citrate carrier (CIC), is indeed downregulated in OsiR (−0.476-fold, adj. *p*-value 1.86 ×10^−4^).

**ACLY**, which catalyzes the ATP-dependent conversion of citrate and CoA to oxaloacetate (OAA) and acetyl-CoA employed in lipogenesis, is also upregulated in OsiR cells (+0.270-fold, adj. *p*-value 4.02 ×10^−6^). Also related to **lipogenesis**, it is worth noting the significant upregulation in OsiR cells of the gene encoding dehydrogenase/reductase member 2 **DHRS2** (+1,056-fold, adj. *p*-value 6,84 ×10^−11^), which belongs to the short-chain dehydrogenases/reductases (SDR) family. These enzymes have essential roles in reprogramming lipid metabolism and redox homeostasis to regulate proliferation, migration, invasion, and drug resistance of cancer cells^128^.

**PDH** is the gatekeeper enzyme in glucose metabolism, directing pyruvate either towards the TCA cycle as acetyl-CoA or toward lactate. Its activity is tightly regulated by pyruvate dehydrogenase kinase (PDK), which phosphorylates and inactivates one of the PDH subunits, and by a corresponding phosphatase that restores PDH activity. Consequently, PDH inhibition or PDK overexpression is expected to promote glycolysis. In OsiR cells, **PDK1**, which inhibits **PDHA1** activity by phosphorylation, is downregulated (−0.311-fold, adj. *p*-value 3.33 ×10^−4^), as is PDHA1 itself (−0.194-fold, adj. *p*-value 3.40 ×10^−3^).

Moreover, the transcriptional stoichiometry of PDH is subtly altered, with PDHA1 being slightly downregulated and PDHB being similarly upregulated (−0.214-fold, adj. *p*-value 2.04 ×10^−4^).

Despite these transcriptional changes, Western blot analysis shows a significant overexpression of PDH protein^78^, and, specifically, of PDHE1 (**Figure 2I**) involved in a atypical decarboxylation step. PDHA1 dysregulation has been implicated in metabolic reprogramming, and PDHA and **PDHB** expression profiles vary across different cancer types and drug resistance models, suggesting diverse mechanisms and functions^129^. PHDB was also found to contribute to drug resistance in lung cancer cells through the promotion of EMT^130^. However, the exact mechanism underlying these adaptations remains unknown. Pyruvate is transported into the mitochondrial matrix via a specific carrier located in the inner mitochondrial membrane, known as the mitochondrial pyruvate carrier (MPC)^131^, where it can be directly converted by pyruvate carboxylase (PC) into OAA, a precursor for several pathways beyond the TCA cycle. Many tumors downregulate MPC to limit pyruvate oxidation^132^. In particular, MPC1 has been identified as an essential factor for sensitizing NSCLC lung cancer lines to PARP inhibition ^133^.

Interestingly, except for the TCA genes and related proteins described above, all the other enzymes of the TCA cycle are equally expressed in Par and OsiR cells, although mutations in ETC complexes (MCI, MCIII, MCIV and ATPase) and functional tests^78^ suggest that in OsiR cells, ETC OXPHOS function is compromised, and mitochondrial energy production is severely impaired.

Thus, mitochondria are likely repurposed from their normal modus operandi to support biosynthetic and anaplerotic processes rather than OXPHOS. Metabolites such as OAA and alpha-ketoglutarate (αKG) have indeed key roles as building blocks of amino acids and, vice versa, serve as entry points of alternative amino acidic-derived carbon sources in cell metabolism. This implies that when the TCA cycle is impaired due to stress and/or mutations, it might not be as critical as previously thought^134^, and cells may adapt by redirecting metabolism to other pathways to maintain functionality^135–137^. An intriguing, challenging hypothesis is that the bioenergetic function of mitochondria is voluntarily “switched off” by the cell in response to specific signaling, reassigning mitochondria the mentioned anaplerotic role.

### 4.1. A special focus on pyruvate, acetaldehyde and acetate: an alternative metabolic route in OsiR cells

The chromatographic analysis evidenced that the exometabolome of Par cancer cells is characterized by higher concentrations of lactate (0.429±0.021 nmol/cell), as reported above, followed by pyruvate (0.022±0.004 nmol/cell), acetaldehyde (0.0019±0.0005 nmol/cell) and no acetate, whose levels are below the instrumental limit of detection (**Table 2**). In contrast, the exometabolome of OsiR cells shows acetate as the main metabolite (1.144±0.219 nmol/cell), followed by lactate (0.606±0.047 nmol/cell), pyruvate (0.113±0.018 nmol/cell) and acetaldehyde (0.0050±0.0009 nmol/cell). Interestingly, lactate (pK_a_=3.1) is responsible for the acid pH of parental ECM; acetate (pK_a_=4.8) is responsible for the acidity of the ECM of OsiR.

While the lactate production could be expected in drug resistant cancer cells because of the enhancement of the Warburg effect, the unexpected high concentrations of pyruvate, acetaldehyde and acetate, and other crucial metabolites, suggest that OsiR cells select specific pathways of glycolysis, TCA and PPP to produce energy, reducing power and building blocks for all the biosynthesis. Considering that hypoxia and acidosis (singly or in combination) cause different responses, both in gene expression and in de novo protein synthesis this may have currently unknown crucial implications in cell survival ^136^.

Many proteins involved in **pyruvate** metabolism are differentially regulated in cancer and normal cells, and many glycolytic enzymes are involved in drug resistance^3^. It is reported that the downregulation of mitochondrial oxidative metabolism (long-term Warburg effect) can be a strategy for escaping cell death^4^. However, cancer cells can reversibly regulate their glucose-induced suppression of respiration and OXPHOS, the so-called **“Crabtree effect”** ^4,6,93,138,139^, although its mechanism is currently unknown. Diaz-Ruiz et al. discussed similarities between the glucose-induced repression of oxidative metabolism in *S. cerevisiae* and the “aerobic glycolysis” of tumor cells^132^, especially in the activity and/or expression pattern of key glycolytic enzymes, finding that specific isoforms expressed in tumor cells have a key role for the induction of drug resistance^3^. As in tumors, in *S. cerevisiae* the pyruvate concentration level is crucial and during fermentation its cytoplasmic metabolism is high and the pyruvate entry into TCA is limited. This last step is finely regulated by the PDH kinase and phosphatase and by the transcriptional regulation of its E3 subunit (lipoamide dehydrogenase). If the transport of pyruvate into mitochondria is lowered and the acetyl-CoA formation is inhibited, the pyruvate concentration level in the cytosol increases. In the case of cancer cells, the TCA downregulation may be a consequence of the compromised mitochondrial respiratory chain, addressing pyruvate toward other metabolic pathways and the anaplerotic role of mitochondria^135–137^.

The role of **acetate** metabolism in cancer cells was recognized as early as 1957^140^ by Weinhouse et al. who demonstrated that acetate can be used as an important respiratory substrate.^69^ It has been reported that acetate can be used as a carbon source for lipid synthesis and energy production, especially when cancer cells are under metabolic stress or in hypoxic conditions ^80^. However, its source and its role in tumor metabolism and in tumor immune evasion remains unclear^141^ and deserves deeper investigations.

In OsiR acetate is the most abundant short-chain fatty acid in the ECM, suggesting that OsiR might rely more on acetate for the biosynthesis of cellular components necessary for survival in the presence of Osimertinib.

The integration of metabolomic, transcriptomic and Western blot data from Par cells and OsiR^78^ allowed us to suggest a specific pathway potentially responsible for the reprogramming of cellular metabolism. Acetate (**Figure 2B** and **Table 2**) was present only in OsiR ECM (1.144±0.219 nmol/cell), and acetaldehyde concentration (**Figure 2G** and **Table 2**) was significantly higher (0.0050±0.0009 nmol/cell *vs* 0.0019±0.0005 nmol/cell in Par). The reasons for the acetaldehyde concentration three order of magnitude lower than acetate could be due both to its partial loss due to the low atmospheric boiling point and to the membrane barriers through which acetaldehyde can diffuse from its site of production. If acetaldehyde is produced in cells due to the imbalanced substrate and cofactor concentrations affecting PDH function, as suggested by Liu et al.^60^, PDH is in fact located in the inner mitochondrial membrane and acetaldehyde can diffuse both in the mitochondrial intermembrane space and in the mitochondrial matrix^142^. Finally, acetaldehyde is effectively decomposed in cells due to its high toxicity, through oxidation to ethanol by ADH (in liver) of CYP2E1^48^, or to acetate by the NADPH-dependent aldehyde dehydrogenases (ALDH family)^143^. No traces of ethanol were found in the exometabolome of Par/OsiR cell lines. Considering the μM concentrations of acetaldehyde measured by Liu et al. in cell lysate models^60^ it seems likely that the upregulation of glycolytic pathway and of ALDHs can produce μM concentration of acetate in the OsiR ECM.

The massive presence of acetate and the increased expression of PDH in OsiR suggests that, along with the glycolysis enhancement, pyruvate follows a specific pathway for the formation of acetate (**Scheme 1**), in agreement with the results found by Liu et al. ^60^ in several human cancer cell lines.

**Scheme 1.**
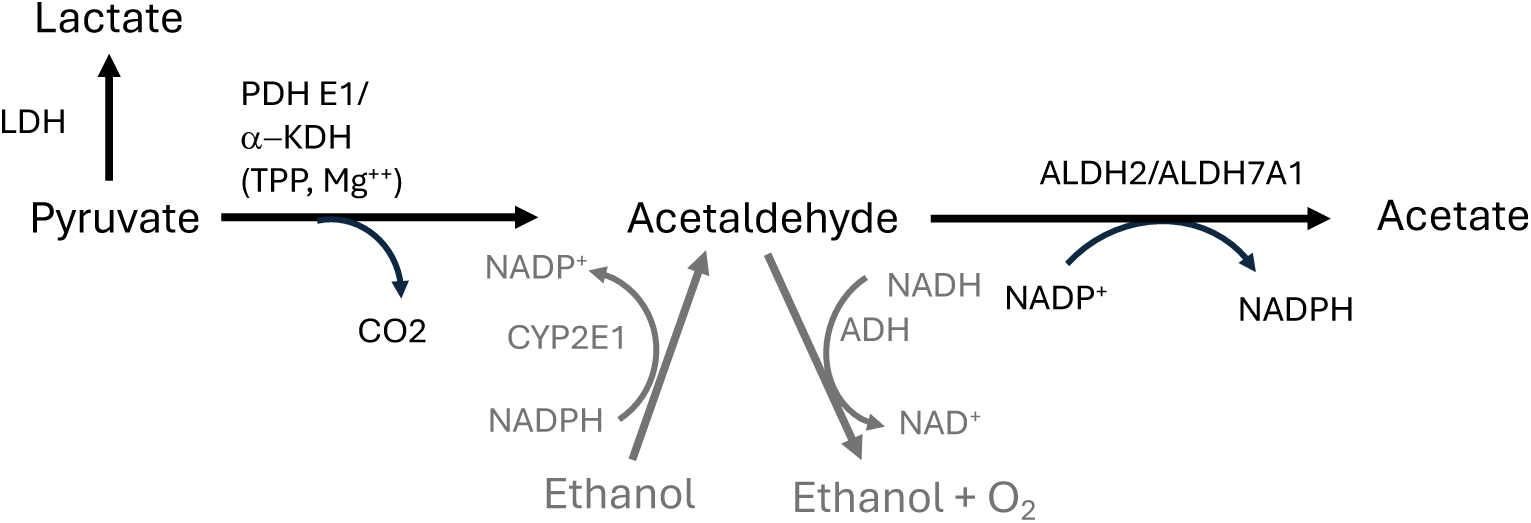
Formation of acetate in OsiR through the pyruvate-acetaldehyde-acetate (PAA) pathway.

To date, 19 ALDH genes have been identified in humans, coding for cytosolic, mitochondrial, and microsomal enzymes in different tissue types ^144–147^. Aldehyde dehydrogenases (ALDHs) have a specific role in the oxidation of acetaldehyde to acetate according to the reaction reported in Scheme 1.^4,10,148–153^ The ALDH family is closely related to the proliferation, migration, invasion and resistance of tumor cells, and different ALDH subtypes are expressed in different tumor cells ^150–152,154^. He et al. have shown by real-time quantitative PCR and Western blotting that in cisplatin-resistant human A549/DDP lung adenocarcinoma cell line the expression of ALDH which was 97.7% vs only 4.3% in parental A549 cells ^151^. They also found that the ALDH inhibitor DEAB was able to significantly reduce cisplatin resistance in A549/DDP cells. In particular, the ALDH1B1 isozyme may play a role in the chemoresistance of these cells; a role for this enzyme was reported in ethanol metabolism with an exclusive preference for NAD^+^ as cofactor and an apparent K_m_ of 55 μM for acetaldehyde, making it the second low-K_m_ (high affinity) ALDH for the metabolism of this substrate^152^.

Yang et al. have found that ALDH1A1 may play a role in gefitinib resistance of lung cancer HCC-827/GR cells ^150^.

In OsiR cells several ALDHs are significantly up- or downregulated (absolute adj. *p*-value >0.3): ALDH2 (+1.240-fold, adj. *p*-value 3.49 ×10^−6^), ALDH7A1 (+0.440-fold, adj. *p*-value 4.46 ×10^−7^), ALDH5A1 (−2.25-fold, adj. *p*-value 5.89 ×10^−8^), ALDH3B1 (−1.22-fold, adj. *p*-value 2.85 ×10^−8^), LDH6A1 (−0.958-fold, adj. *p*-value 1.47 ×10^−6^), ALDH3A2 (−0.305-fold, adj. *p*-value 7.89 ×10^−5^). These data show a clear rearrangement of the acetaldehyde metabolism. In liver it is known that two major isoforms of ALDH, cytosolic and mitochondrial, can be distinguished by their electrophoretic mobilities and kinetic properties^155^.

In OsiR cells, ALDH2 (1.2-fold, adj. *p*-value =3.5 ×10^−6^) and ALDH7A1 (0.44-fold, adj. *p*-value =6.6 ×10^−7^) are upregulated both at mRNA and protein level^78^ with respect to Par cell lines. In humans ALDH2 is especially efficient on acetaldehyde compared to ALDH1^156^, and it has specific functions against oxidative stress^157^. ALDH7A1 is found both in the cytosol and the mitochondria and it performs NAD(Par)^+^-dependent oxidation of aldehydes ^158,159^.

ALDH2 encodes a mitochondrial matrix protein that is constitutively expressed in a variety of tissues^160^. ALDH2 is the primary enzyme involved in the oxidation of acetaldehyde (K_m_ < 1 μM) during ethanol metabolism ^161^. Other authors found that human ALDH2 exhibits the highest affinity for acetaldehyde with K_m_ = 0.59–0.27 μM ^156^ (0.2 μM in Klyosov et al.^161^). Mitochondrial aldehyde dehydrogenase ALDH2 has been reported to reduce ischemic damage in an experimental myocardial infarction model and its activity is redox-sensitive ^162,163^. Upregulation of ALDH2 expression in NSCLC cells is highly associated with resistance to Paclitaxel in vitro and in vivo ^164^, and Disulfiram in combination with copper reverses microtubule inhibitor resistance in cancer cells by suppressing ALDH2 expression in NSCLC cells^164^. Moreb at al. found that overexpression of ALDH1A2 or ALDH2 in K562 leukemia and H1299 lung cancer cell lines exhibited higher cell proliferation rates, higher clonal efficiency, and increased drug resistance to 4-hydroperoxycyclophosphamide and doxorubicin ^165^.

ALDH7A1 is involved in the pipecolic acid pathway of lysine catabolism, which can regulate osmotic pressure^158^, being involved in the cellular defense against hyperosmotic stress and in attenuating reactive aldehyde and oxidative-stress-induced cytotoxicity^158,159^. **Lysine** (**Figure 3A**) and **pipecolic acid** (**Figure 3B**) are indeed significantly higher in OsiR cells.

ALDH7A1 is the only ALDH found in the cytosol, nucleus, and mitochondria that is differentially expressed in a tissue-specific manner in mice ^159^.

Human ALDH7A1 showed high specificity for the medium-chain saturated aldehydes that are known to be potentially toxic lipid peroxidation products ^166^. Little kinetic information exists regarding mammalian ALDH7A1 activity towards other substrates^160^.

ALDH isoforms (ALDH5A1, ALDH3B1, ALDH6A1 and ALDH3A2) downregulated in OsiR are instead unrelated to acetaldehyde oxidation. ALDH5A1 encodes a mitochondrial NAD^+^-dependent succinic semialdehyde dehydrogenase. ALDH3B1 is cytosolic, NAD^+^/NADP^+^-dependent and catalytically active toward C16 aldehydes derived from lipid peroxidation, suggesting a potential role against oxidative stress ^149,167,168^, as well as ALDH3A2 ^168^. ALDH3B1 demonstrated high affinity for hexanal (K_m_=62 μM), octanal (K_m_=8 μM), 4-hydroxy-2-nonenal (K_m_=52 μM), and benzaldehyde (K_m_=46 μM), but low affinity toward acetaldehyde (K_m_=23.3 mM) ^149^. The ALDH6A1-encoded protein is a mitochondrial methylmalonate semialdehyde dehydrogenase that plays a role in the valine and pyrimidine catabolic pathways. ALDH6A1catalyzes the irreversible oxidative decarboxylation of malonate and methylmalonate semialdehydes to acetyl- and propionyl-CoA and it is downregulated in gastric cancer ^169^.

ALDH inhibitors have indeed been proposed for cancer therapeutics and to overcome chemotherapy resistance^154,170–172^. However, the role of ALDH in cancer cell metabolism is currently not fully investigated.

*Why should this kind of cellular reprogramming be activated under life-threatening conditions like chemotherapy?* The meaning of the pathway outlined above (**Scheme 1**) has not previously been reported.

In their work Liu et al. conclude that they “have by no means defined the general biological role of these reactions”^60^. We hypothesize here that the high concentration of acetate is a consequence of the activation of an alternative route to generate ATP, which is efficient not in yield but in speed, through glycolysis and reducing power (NADPH) production under critical conditions. In other words, we propose here that ALDH2 and ALDH7A1 have a specific role in the production of NADPH (**Scheme 1**), and that this pathway could be specifically switched on to allow the survival of cells developing drug resistance.

This interpretation fits the transcriptomics and the data available for OsiR: (i) the upregulation of glycolytic-pathway proteins (Warburg effect), (ii) the down regulation of ox-PPP and, thus, the PPP-mediated NADPH production, (iii) the enhanced biosynthesis of lipids in OsiR^78^, (iv) impaired OXPHOS and mitochondrial function related to energy production^78^.

Bose et al. also reported that under conditions such as mitochondrial dysfunction, hypoxia, limited metabolic resources, and in tumor environments, the usual pathway converting pyruvate to acetyl-CoA can change, and cells start using acetate as a major energy source ^61^. However, beyond its role as a carbon source and as a promoter of immune evasion^141^, the PAA pathway also has the crucial role of producing NADPH and of readdressing pyruvate toward acetate to supply cytoplasmic acetyl-CoA for lipid biosynthesis.

Acetyl-CoA is also generated in the mitochondria from pyruvate by PDH and in the cytosol from citrate by ACLY, which is indeed upregulated in OsiR, as reported above. Both processes are coupled to the TCA cycle. Moreover, **ACSS1** and **ACSS2** catalyze the conversion of endogenous acetate to acetyl-CoA in the mitochondria and cytosol, respectively, which is crucial for lipogenesis^141^. In OsiR cells mitochondrial ACSS1 is significantly more expressed than in Par cells (**Figure 2J**). ACSS2 has been reported to be crucial for sustaining acetyl-CoA pools and supporting cell proliferation in metabolically limited environments, such as during mitochondrial dysfunction or ATP citrate lyase (ACLY) deficiency^173,174^. This highlights the ACSS importance in maintaining cellular metabolism under stress conditions. Moreover, in OsiR cells, the downregulation of isocitrate dehydrogenase (IDH1 and IDH2) suggests a reorientation of citrate metabolism towards feeding the acetyl-CoA pool.

While NADPH is primarily used in anabolic reactions such as fatty acid biosynthesis, NADH, in the presence of fully or partially preserved mitochondrial function, is more involved in catabolic reactions and energy production^175^. NADH/NADPH balance in cells is in fact regulated by nicotinamide-nucleotide transhydrogenase (NNT), a membrane-bound protein facilitating the reduction of NADP^+^ by NADH through proton translocation. In eukaryotic cells, NNT in mitochondria translocates protons from the intermembrane space to the matrix, and in the endoplasmic reticulum (ER), it moves protons from the inner ER to the cytosol. NNT activity impacts redox status, biosynthesis, detoxification (via glutathione/thioredoxin systems), and apoptosis and it is involved in various pathological conditions, including cancer cell proliferation.^176,177^. However, the role of NNT in NSCLC remains debated^176,178–182^. In both Par and OsiR cells, NNT transcription and protein levels are unchanged (data not shown), likely because NADPH/NADH levels are similar, despite being produced through different pathways (ox-PPP in Par cells, and PAA in OsiR cells).

### 4.2. Anaplerotic Flux and Alternative Pathways

Beyond glycolysis, OsiR cells also activate anaplerotic pathways to replenish intermediates in glycolysis, the PPP and the TCA cycle, supporting biosynthetic needs under stress. For instance, 3-PG leads to the formation of serine (**Figure 3C**). Histidine (**Figure 3D**) has increased levels in OsiR. It is biosynthesized from ribose-5-phosphate and it is linked to *de novo* purine biosynthesis. Histidine is reported to contribute to chemoresistance in some cancers^183^. Glycine (**Figure 3E**) and cystine (**Figure 3F**), the oxidized species of cysteine, derive from serine. Shifts in **serine and glycine** metabolism support one-carbon metabolism, which is crucial for maintaining nucleotide synthesis and TCA cycle function. Additionally, **glycine** serves as a substrate for the synthesis of **glutathione**, an important antioxidant that helps mitigate oxidative stress, a hallmark of drug resistance.

Pyruvate contributes to the synthesis of alanine (**Figure 3G**). Additionally, intermediates from the TCA cycle, such as OAA, produce aspartate (**Figure 3H**) and asparagine (**Figure 3I**), while α-KG gives rise to arginine (**Figure 3J**).

Our analysis revealed significant changes in **amino acid metabolism**, including the increase of branched-chain amino acids (**valine** and **isoleucine, Figure 3K** and **3L**, respectively). These pathways are crucial for maintaining metabolic flexibility and survival in the face of drug-induced stress.

These alterations contribute to the replenishment of key TCA cycle intermediates like α-ketoglutarate, which helps maintain TCA cycle activity when mitochondrial function is compromised. Valine and isoleucine are converted into intermediates like propionyl-CoA, which can enter the TCA cycle. **Phenylalanine** (**Figure 3M**) and **tyrosine** (**Figure 3N**) serve as precursors for α-ketoglutarate, supporting the anaplerosis of the TCA cycle. Although **tryptophan** (**Figure 3O**) is primarily involved in **serotonin** (**Figure 3P**) and niacin biosynthesis, it indirectly supports TCA cycle function, by contributing to NAD^+^ production, primarily through kynurenine metabolism. Elevated **kynurenine** levels **(Figure 3Q),** along with 2-picolinic acid **(Figure 3R)**, a metabolite of tryptophan via the kynurenine pathway, were observed in OsiR cells. This suggests that tryptophan catabolism via kynurenine is critical role in modulating immune responses in the tumor microenvironment (TME) and maintaining redox balance during stress. Moreover, the kynurenine pathway helps regulate cellular responses to oxidative stress, which could contribute to immune evasion and drug resistance in OsiR cells.

The increase in oxidized amino acids (such as **methionine sulfoxide**, **Figure 3S**, a marker of amino acid oxidation) and **cystine** indicates an altered redox status in OsiR cells. Additionally, the increase in **2-oxovaleric acid** (**Figure 3T)** and **3-methyl-2-oxopentanoic acid** and **4-methyl-2-oxopentanoic acid** (or α-ketoisocaproic acid, A-KIC) (**Figure 3U** and **3V)**, which are short-chain keto acids, suggests active metabolism in OsiR cells. These secondary metabolites, although not essential for basic cellular functions, may act as defense or signaling molecules^184^. **Hippuric acid (Figure 3W)**, a major conjugate of aromatic amino acids, was also found to be elevated in OsiR cells, possibly indicating an enhanced detoxification process to mitigate reactive metabolites, though its role is complex^185^.

These changes suggest that OsiR cells are relying on a combination of glycolytic and anaplerotic pathways to bypass the limitations imposed by impaired mitochondria, maintaining cellular function and survival despite the stress from therapy.

### Nucleotide Metabolism

Metabolic profiling revealed increased levels of **adenine**, **nicotinic acid**, and **N-methyl nicotinamide** (**Figure 3X, 3Y, 3Z,** respectively), along with reduced **nicotinamide** levels (**Figure 3AA**), indicating a shift in nucleotide metabolism. This shift supports cell survival under stress conditions by adjusting NAD^+^ levels, an essential cofactor for redox reactions.

Interestingly, **alcohol dehydrogenase 5 (ADH5),** a NADP^+^-dependent enzyme, is upregulated in OsiR cells (+0.299-fold, adj. *p*-value =4.13 ×10^−5^). It catalyzes the oxidation of S-hydroxymethylglutathione to GSH and formate, which is crucial for ***de novo* synthesis of purines**^186^. Formate, derived from the breakdown of folate derivatives ^187^, is also essential for DNA and ATP synthesis, supporting cell growth and replication^188^. Moreover, folate metabolism is also one of **NADPH production pathways**^189^ that can compensate for the downregulated oxPPP ^190,191^.

In mitochondria, along with two enzymes linked to one-carbon (1C) metabolism (the aldehyde DH, ALDH1L2, and methylenetetrahydrofolate dehydrogenase, MTHFD2) that produce NADPH and formate, NADPH is produced also by IDH2, which is upregulated in OsiR, as mentioned above^190^. In the cytosol the reverse reactions catalyzed by ALDH1L1 and MTHFD1 consume NADPH. The canonical function of folate metabolism is to provide activated one-carbon units to enable synthesis of thymidine, purines, and methionine^191^. In the mitochondria, NADPH would provide reducing power to neutralize ROS. In the cytosol, NADPH is produced by G6PDH (the rate-limiting step of the PPP), by IDH1, and malic enzyme that converts malate into pyruvate; interestingly, in OsiR ME2 and ME3, i.e. the genes encoding mitochondrial malic enzyme isoforms, have a tendency to upregulation (+0.110-fold, adj. *p*-value =1.05 ×10^−2^ and +0.217-fold, adj. *p*-value =3.60 ×10^−2^, respectively). In the cytosol, NADPH is mainly used for reductive biosynthesis of lipids, cholesterol, and steroids, and reducing power to neutralize ROS. In the ER, NADPH provides reducing power for disulfide isomerization and contributes to steroid activation^192^.

In OsiR, H6PDH and its cytoplasmic homolog G6PDH, whose reactions are considered the major source of NADPH, are downregulated, as reported above. In NSCLC with acquired drug resistance, cytosolic ME and G6PDH have been reported as key regulators of redox balance^193,194^

**Nicotinic acid** and **adenine** levels, which are significantly increased, and **nicotinamide** (vitamin B3), which is decreased in OsiR cells, are precursors of the coenzymes nicotinamide-adenine dinucleotide (NAD^+^) and nicotinamide-adenine dinucleotide phosphate (NADP^+^). NAD^+^ synthesis occurs either de novo from amino acids, or through a *salvage pathway* from nicotinamide which is only typically found in mammals. The *de novo* pathway includes the formation of quinolinic acid. In OsiR cells the levels of quinolinic acid are significantly lower (and below the instrumental detection limit) than those observed in parental cells (**Figure 3AB**). Quinolinic acid is a downstream product of the kynurenine pathway, which metabolizes tryptophan for the NAD^+^ synthesis, and it is linked to tumor growth, progression, and resistance to therapy. The use of quinolinic acid for NAD^+^ synthesis is known to prevent apoptosis, increase resistance to oxidative stress induced by radiochemotherapy, conferring a poorer prognosis in gliomas^195^.

The metabolite changes suggests an **upregulation of the NAD^+^ salvage pathway**, crucial for maintaining redox balance and energy metabolism under drug-induced stress (http://pathbank.org/view/SMP0000048)^80^.

**Higher adenine levels** may be due to some recycling process, from 5’-thiomethyl-adenosine or from 2’-deoxyadenosine. Purine nucleotides can be provided by *de novo* or *salvage pathways*, but the increase in adenine is indicative of a surplus deriving from other pathways. Interestingly, the AOX1 gene encoding for aldehyde oxidase is significantly upregulated in OsiR cell (+1.911-fold, adj. *p*-value 4.84 ×10^−8^). AOX1 produces hydrogen peroxide and, under certain conditions, can catalyze the formation of superoxide. AOX1 is involved in the NAD^+^/NADP^+^ salvage pathway. In colorectal cancer it is known to promote cancer progression by upregulating CD133 expression^196^.

Antiporters of adenine nucleotides are also upregulated in OsiR cells (SLC25A23 and SLC25A24, i.e. −0.392-fold, adj. *p*-value 3.50 ×10^−6^ and +0.444-fold, adj. *p*-value 4.01 ×10^−6^), respectively).

OsiR cells also show a significant **reduction in nicotinamide** and the corresponding increase of **1-methylnicotinamide** (1-MNA), indicating a potential rerouting of NAD^+^ biosynthesis pathways (salvage pathway) in response to altered redox states. This matches the upregulation of Nicotinamide N-methyltransferase (**NNMT)**, which transfers a methyl group from S-adenosyl methionine (SAM) to nicotinamide, producing S-adenosyl-L-homocysteine (SAH) and 1-methylnicotinamide (1-MNA). High NNMT expression alters SAM levels, affecting NAD^+^-dependent redox reactions and signaling pathways ^197^. NNMT plays a significant role in lung cancer progression ^198^ by inhibiting DNA repair and promoting cell cycle progression. Its product, 1-MNA, inhibits autophagy and apoptosis ^199–201^. Overexpression of NNMT is linked to acquired resistance to EGFR-TKI^202^, and histone and DNA methylation, leading to changes in gene expression that support cancer cell survival and proliferation^203^. NNMT supports NAD^+^-consuming enzymes, crucial for maintaining cellular redox balance and energy metabolism, and can promote the Warburg effect, where cancer cells rely more on glycolysis for energy production, even in the presence of oxygen^204^. Interestingly, in OsiR cells the genes related to the transport of S-adenosyl-methionine (SAM) and S-adenosyl-homocysteine (SAH) into mitochondria are upregulated (SLC25A26 and SLC25A28, i.e. +0.202-fold, adj. *p*-value 7.44 ×10^−3^ and +0.201-fold, adj. *p*-value 3.45 ×10^−2^), respectively).

### 4.3. Neurotransmitters

L-DOPA or l-3,4-dihydroxyphenylalanine is the precursor to the neurotransmitters dopamine, norepinephrine (noradrenaline), and epinephrine (adrenaline), collectively known as catecholamines. L-DOPA, is implicated in the regulation of tumor growth^205^ and may impact the pathogenesis of glioma by influencing angiogenesis, vasculogenesis, apoptosis, and autophagy^206^.

Neurotransmitter metabolism was impacted is osimertinib resistance, with decreased serotonin (**Figure 3P**) and increased DOPA levels in OsiR cells (**Figure 3AC**), pointing to broader metabolic adaptations that may influence cellular signaling and stress responses.

### 4.4. Acidity-related transporters

The presence of acetate and lactate in OsiR cells makes the exometabolome to become acidic, and many genes related to the control of cell acidity are upregulated with respect to Par cells.

SLC4A7 (+0.605-fold, adj. *p*-value 4.61 ×10^−8^) functions as an electroneutral Na^+^/HCO_3_^−^ cotransporter, playing a crucial role in acid extrusion after cellular acidification^207^. It also provides cellular bicarbonate for *de novo* purine and pyrimidine synthesis, acting as a key mediator of nucleotide synthesis downstream of mTORC1 signaling in proliferating cells.

Similarly, SLC2A13 (+0.859-fold, adj. *p*-value 3.08 ×10^−5^) encodes a H^+^ myo-inositol cotransporter, which is also identified as a potential marker for cancer stem cells in oral squamous cell carcinoma^208^, and SLC9A1 gene (+0.407-fold, adj. *p*-value 3.94 ×10^−5^), encoding the sodium-hydrogen exchanger 1 (NHE-1), are upregulated in OsiR cells. NHE-1 primarily regulates intracellular pH by exchanging intracellular hydrogen ions for extracellular sodium ions (Na⁺), which is crucial for maintaining cellular homeostasis^209^.

Glucose transporters are generally downregulated is OsiR (SLC2A12, SLC2A3, and SLC2A8, i.e. −0.605-fold, adj. *p*-value 3.75 ×10^−3^, −0.215-fold, adj. *p*-value 3.89 ×10^−2^, and −0.529-fold, adj. *p*-value 3.68 ×10^−5^, respectively), yet the sucrose-proton symporter activity is supported by the upregulation of SLC45A3 (+0.456-fold, adj. *p*-value 1.09 ×10^−4^) and SLC45A4 (+0.197-fold, adj. *p*-value 4.42 ×10^−2^).

## 5. Conclusions

The exometabolome analysis of OsiR EGFR-mutant NSCLC compared to the EGFR-mutant H1975 cell line revealed distinct metabolic profiles. Par cells exhibited higher extracellular concentrations of lactate, followed by pyruvate and acetate (with acetate levels below the instrumental limit of detection). In contrast, in OsiR cells, targeted characterization of the exometabolome unexpectedly identified acetate as the predominant source of extracellular acidity, followed by lactate and pyruvate. These finding suggest that resistant cells may rely more on glycolysis (Warburg effect), as evidenced by increased glycolytic activity, overexpression of specific glycolytic enzymes, and activation of alternative metabolic pathways supporting survival in the presence of osimertinib. The markedly elevated acetate levels in the OsiR exometabolome, alongside and other distinct metabolic features in OsiR cells metabolites suggest the occurrence of a specific resistance-related metabolic switch. Preliminary transcriptomics, mitochondrial genomic, and imaging analyses of OsiR cells^78^ have shown profound alterations in mitochondrial function and morphology, including impaired ETC, the presence of mtDNA mutations, and the downregulation of several mitochondrial genes, all consistent with compromised OXPHOS.

We hypothesize that the overproduction of acetate as well as the increased pyruvate levels in OsiR cells are due to the activation of an unconventional metabolic route: the pyruvate-acetaldehyde-acetate pathway. This pathway may serve as the main source of NADPH, supporting lipid biosynthesis and oxidative stress management in resistant cells. Consistent with this hypothesis, we observed the upregulation of ALDH2 and ALDH7A1. The upregulation of ALDH2 and ALDH7A1, enzymes that catalyze the NADP^+^-dependent oxidation of acetaldehyde to acetate, generating NADPH.

Interestingly, while the enzymes of the oxidative branch of the PPP (G6PDH, H6PDH, PGLS) are downregulated in OsiR cells, those involved in the non-oxidative branch of the PPP are equally expressed in Par and OsiR cells. This suggests that both NADPH and the non-ox-PPP metabolites required for *de novo* lipid biosynthesis are replenished through different mechanisms. Specifically, in Par cells NADPH mainly derives from the ox-PPP, while metabolites from the ox-PPP; in OsiR cells, NADPH is generated via the PAA pathway, while metabolites are produced by the glycolytic intermediates (F6P and G3P) via the “shunt” enzyme TKT. Both NADPH and acetate are crucial for acetyl-CoA production and *de novo* lipid biosynthesis.

Furthermore, the TCA cycle likely plays an anaplerotic role in OsiR cells, providing precursors for amino acids, nucleotides, vitamin, and cofactor biosynthesis. Western blot analyses showed increase expression of two enzyme that are crucial for mitochondrial metabolism (PDH and SDH), in OsiR cells, supporting this hypothesis.

Overall, this metabolic reprogramming, together with the upregulation of stress response genes^78^, highlights a comprehensive cellular adaptation strategy. It illustrates how OsiR cells maintain viability despite a profoundly compromised mitochondrial “power house” function, while mitochondria continue to actively provide key metabolites necessary for biosynthetic processes.

## FUNDER INFORMATION DECLARED

AIRC, Investigator Grant 2021 (ID 25734); PNNR, MCNT1-2023-12377671 Grant; Pisa Foundation Grant, ELMO Fondazione Pisana per la Scienza, FPS Grant 2024; European Union - Next Generation EU, Mission 4 Component 2, project “Strengthening BBMRL.it”; NUS, A-0001263-00-00

